# Status of susceptibility of the visceral leishmaniasis vector, *Phlebotomus argentipes* (Diptera: Psychodidae: Phlebotominae), to insecticides used for vector control in Nepal

**DOI:** 10.1101/2022.03.07.483225

**Authors:** Lalita Roy, Surendra Uranw, Kristien Cloots, Tom Smekens, Usha Kiran, Uttam Raj Pyakurel, Murari Lal Das, Rajpal S. Yadav, Wim Van Bortel

## Abstract

**Background:** Visceral leishmaniasis (VL) is targeted for elimination as a public health problem in Nepal by 2023. For nearly three decades, the core vector control intervention in Nepal has been indoor residual spraying (IRS) with pyrethroids. Considering the long-term use of pyrethroids and possible development of resistance of the vector *Phlebotomus argentipes* sand flies, we monitored susceptibility status of their field populations to the insecticides of different classes, in villages with and without IRS activities in recent years.

**Methodology/Principal findings:** Sand flies were collected from villages with and without IRS in five VL endemic districts from August 2019 to November 2020. The WHO susceptibility test procedure was adopted using filter papers impregnated at the discriminating concentrations of insecticides of the following classes: pyrethroids (alpha-cypermethrin 0.05%, deltamethrin 0.05% and lambda-cyhalothrin 0.05%), carbamates (bendiocarb 0.1%) and organophosphates (malathion 5%). Pyrethroid resistance intensity bioassays with papers impregnated with 5× of the discriminating concentrations, piperonyl butoxide (PBO) synergist-pyrethroid bioassays and DDT cross resistance bioassays were also performed. In the IRS villages, the vector sand flies were resistant (mortality rate <90%) to alpha-cypermethrin and possibly resistant (mortality rate 90–97%) to deltamethrin and lambda-cyhalothrin, while susceptibility to these insecticides was variable in the non-IRS villages. The vector was fully susceptible to bendiocarb and malathion in all villages. A delayed knockdown time (KDT_50_) with pyrethroids was observed in all villages. The pyrethroid resistance intensity was low, and the susceptibility improved at 5× of the discriminating concentrations. Enhanced pyrethroid susceptibility after pre-exposure to PBO and the DDT-pyrethroid cross-resistance were evident.

**Conclusions/Significance:** Our investigation showed that *P. argentipes* sand flies have emerged with pyrethroid resistance, suggesting the need to switch to alternative classes of insecticides such as organophosphates for IRS. We strongly recommend for the regular and systematic monitoring of insecticide resistance in sand flies to optimize the efficiency of vector control interventions to sustain VL elimination efforts in Nepal.

**Author summary:** Visceral leishmaniasis (VL), transmitted by *P. argentipes* sand flies, is endemic in South-East Asian countries such as Bangladesh, India and Nepal, and is on the verge of elimination as a public health problem in Nepal by 2023. As part of the WHO Global Vector Control Response, entomological surveillance including insecticide resistance monitoring is one of the four main pillars of this strategy. In the early 1990s, the historical use of DDT for sand fly vector control was replaced with deltamethrin or alpha-cypermethrin, which have now been in use for almost three decades in Nepal. Suspecting that this long-term use of pyrethroids might have selected resistance in sand fly populations which would jeopardize control efforts, we conducted the first comprehensive survey to generate contemporary evidence of insecticide resistance in Nepal. For this, we performed WHO susceptibility tests in five VL endemic districts and found strong evidence of pyrethroid resistance in vector populations from the areas receiving IRS. Resistance mechanisms involved would probably be *kdr* mutations and monooxygenase. This study also endorses regular insecticide resistance monitoring to inform evidence-based decisions on insecticide use for vector control and to maintain the effectiveness of vector control measures as a core intervention in the fight against VL.

## Introduction

Indoor residual spraying (IRS) is the core vector control intervention for elimination of visceral leishmaniasis (VL) in South-East Asia, where about 200 million people are at risk of the disease [1, 2]. VL, also called kala-azar in the Indian subcontinent, is a vector-borne neglected tropical disease which is fatal if not treated timely, and mostly affects the lower socio-economic classes of the communities [3]. The causative agent of the disease is the protozoan parasite *Leishmania donovani* Laveran & Mesnil, 1903 (Kinetoplastea: Trypanosomatida) and is transmitted between human hosts by female sand flies of the species *Phlebotomus argentipes* Annandale & Brunetti, 1908 (Diptera: Psychodidae: Phlebotominae) [4]. IRS has shown to reduce the *P. argentipes* density in Nepal [5–7], and is a key intervention in the country’s VL elimination strategy. Since 1992, mostly synthetic pyrethroids i.e. alpha-cypermethrin, deltamethrin or lambda-cyhalothrin have been sprayed in houses [8], after the use of dichloro-diphenyl-trichloro-ethane (DDT) was banned in the country due to its health and environmental hazards [9]. The choice of synthetic pyrethroids over other classes of insecticides for IRS was based on the combination of low cost and high efficacy of these insecticides [2]. However, there are concerns that the use of a single class of insecticides for a long period of time might have potentially selected resistance in sand fly vector and could undermine effectiveness of vector control in Nepal.

At present, little information is available on the current susceptibility status of *P. argentipes* to the pyrethroid insecticides used for IRS in Nepal. Previously reported DDT resistance in *P. argentipes* populations [10] provided a clue for possible cross-resistance to pyrethroids due to involvement of knockdown resistance (kdr) mutations in a common *para-type voltage-gated sodium channel (vgsc)* target site gene in nerve cells [11]. Early signs of pyrethroid resistance were observed in the Eastern part of Nepal and also in some Indian areas bordering the country [11, 12].

Systematic monitoring of insecticide resistance is considered a key component of VL surveillance. The choice for an appropriate insecticide needs to be dynamic and adopted to the changing insecticide resistance status of the vectors if IRS interventions are to be successful [13]. Besides monitoring of resistance to the currently used pyrethroids, it is therefore equally essential to understand the underlying resistance mechanisms and assess the susceptibility status against insecticides of alternative classes with unrelated modes of action, in order to select the most efficacious insecticides, and allow for informed decision-making.

With this study, we aimed to provide an update on the susceptibility status of wild caught *P. argentipes* sand flies to pyrethroid insecticides currently used for IRS as well as to insecticides from two alternative classes with unrelated modes of action (i.e., carbamate and organophosphate) that have never been used for vector control in Nepal. Additionally, we assessed the strength of resistance in pyrethroid resistant phenotypes of sand flies with papers impregnated with five times (5×) the discriminating concentrations of pyrethroids, i.e., alpha-cypermethrin, deltamethrin and lambda-cyhalothrin. We also explored possible underlying metabolic resistance mechanisms by piperonyl butoxide (PBO) synergist-pyrethroid bioassays, and by cross-resistance studies with DDT to ascertain the involvement of *kdr* mutations in the pyrethroid resistant phenotypes.

## Methods

### Study sites

The study was carried out in five VL endemic districts in Eastern and Central Nepal, namely Morang, Saptari, Siraha, Mahottari and Sarlahi (Fig 1). These districts are located at 26°23’-26°55’ N latitude and 85°37′–87°21′ E longitude in the Southern lowland plains of Nepal, which is also known as the ‘Terai’ region. The area has a tropical savanna climate with a mean annual temperature of 19–31 °C and a mean annual rainfall of 1600–2500 mm [14, 15]. These districts were selected based on (i) a high average annual VL incidence reported in the years 2016–2018 and (ii) accessibility by road. In each district, two villages were selected; one intervention village, which received regular or intermittent IRS during the last three or more years due to the presence of VL cases (subsequently called IRS village), and one non-intervention or control village, which did not receive IRS in the three years prior to 2019 as no VL case was reported (subsequently called non-IRS village).

**Fig 1.**
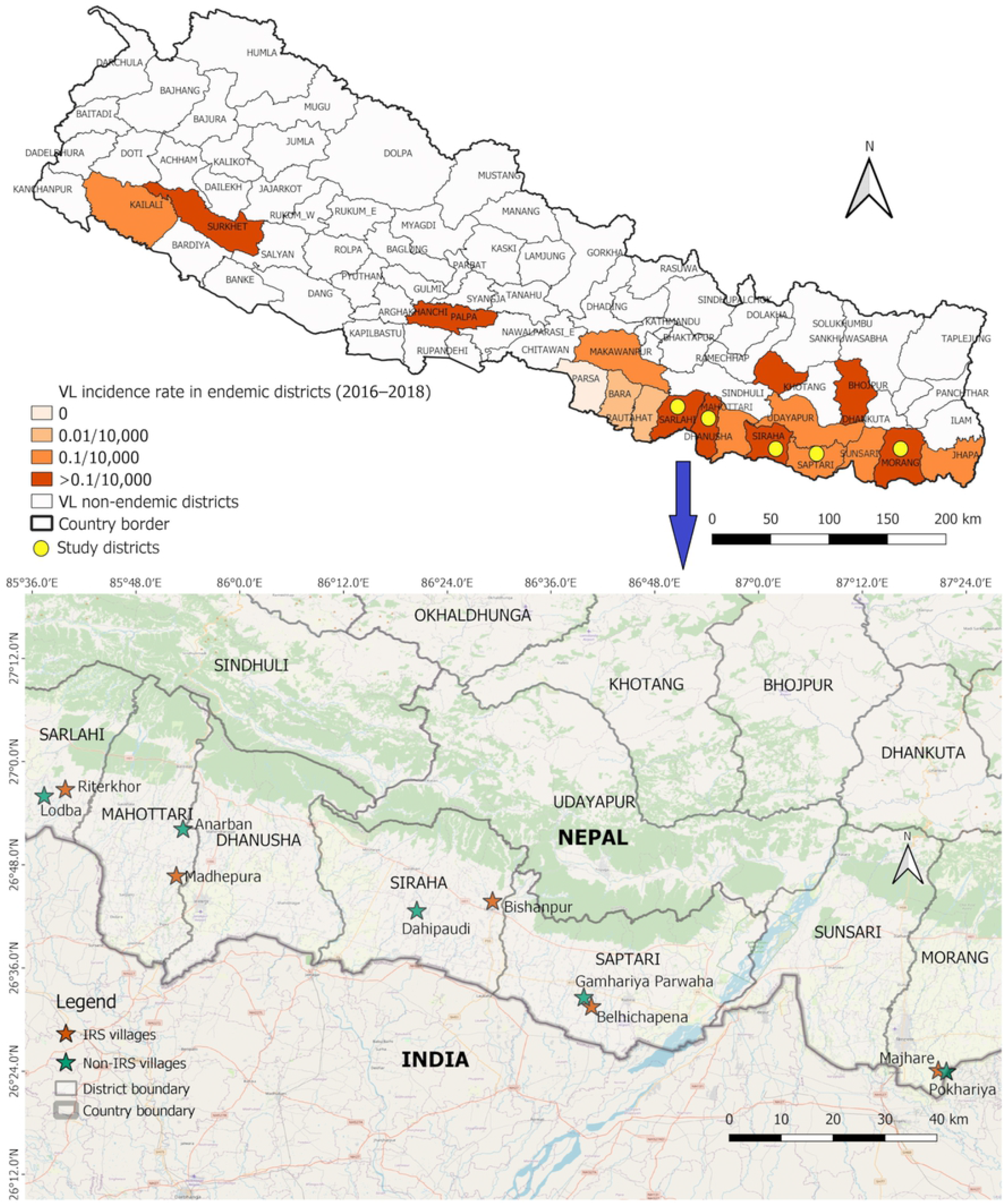
Map of Nepal showing visceral leishmaniasis endemicity (top) and the locations of the study districts and villages (bottom). The IRS villages received spray during three years prior to the start of the study (2016–2018); the non-IRS villages did not receive spray during the same period. The map was produced with QGIS (version 3.10) with shapefiles downloaded from Survey Department (dos.gov.np), Ministry of Land Management, Cooperatives and Poverty Alleviation, Government of Nepal.

### Sand fly collection

Sand flies resting in human dwellings, cattle sheds and dwellings shared by both human and cattle (mixed dwellings) in the selected study villages were collected from August 2019 to November 2020. Collections were done in the early morning between 04:30 and 06:30 hours using a mouth aspirator and a torch. About 5–10 households per village were visited to collect enough sand flies to perform at least one set of bioassays in a day. A temporary laboratory was set up in each village to avoid long distance transportation, which can stress or damage the captured sand flies.

### Bioassays

The WHO bioassay test kit and insecticide impregnated papers were obtained from Universiti Sains Malaysia, Penang, Malaysia. The susceptibility tests with all insecticides and intensity bioassays with pyrethroids were carried out in two phases following the standard WHO susceptibility test procedure [16] (Fig 2). The usual test procedures for mosquitoes were adopted to sand flies. In Phase I of the study, baseline susceptibility status of wild caught female *P. argentipes* was assessed in all the selected villages with filter papers impregnated at the discriminating concentrations of insecticides of the following classes: pyrethroids (alpha-cypermethrin 0.05%, deltamethrin 0.05%, and lambda-cyhalothrin 0.05%), carbamates (bendiocarb 0.1%) and organophosphates (malathion 5%). Silicone oil and olive oil impregnated papers were used as control papers for pyrethroid and organophosphate/carbamate bioassays respectively.

**Fig 2.**
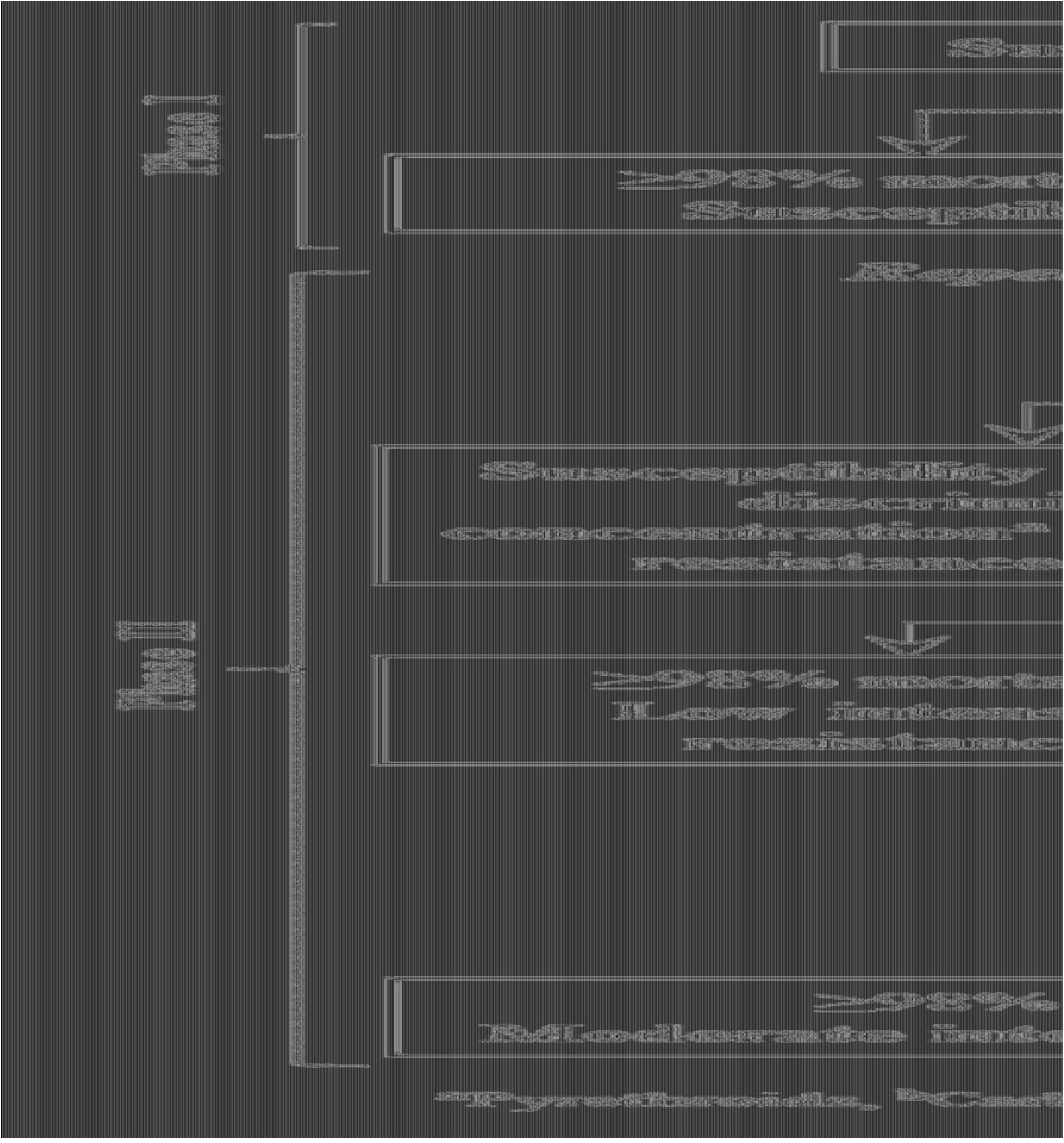
A graphic representation of susceptibility test sequence and probable outcomes for insecticide resistance monitoring in *Phlebotomus argentipes* wild populations. (adopted from the WHO test procedures [16]).

Before exposure, the sand flies were acclimatized for one hour in holding tubes lined with insecticide free clean white papers, each tube holding 20–25 sand flies. They were then transferred from the holding tubes to the exposure tubes lined with insecticide impregnated papers. Each test for a given insecticide impregnated paper comprised of five replicates of 20–25 sand flies, while each control test in parallel comprised of one control tube with a similar number of sand flies. In case of insufficient numbers of sand flies collected i.e., <120 sand flies per set, the test was repeated on subsequent days until 100 or more sand flies had been included in the five replicates, in addition to one control group for each batch of test.

The exposure time was fixed to 60 minutes. Knockdown of the sand flies was recorded 10, 15, 20, 30, 40, 50 and 60 minutes after start of the exposure with insecticides. After 1h of the exposure, the sand flies were transferred back to the holding tubes, which were kept standing upright with the screen end up, provided with 10% glucose-soaked cotton wool pads on the mesh screen, and placed in an area with diffused illumination for the next 24h recovery period. On hot and dry days, the holding tubes were placed in a container box covered with wet towels to stabilize the ambient temperature below 30°C and the relative humidity at 80% ± 10%. The numbers of dead and alive sand flies from each of the exposure and control holding tubes were counted and recorded on a pre-printed form at the end of the recovery period. The species of the sand flies was identified after completing tests with the help of a stereomicroscope, regional taxonomic keys and resources [17–19]. The physiological stages of the sand flies were also recorded. The following criteria were used to classify phenotypic resistance frequency in wild sand flies: fully susceptible (mortality rate ≥98%), possible resistance (mortality rate 90–97%) and confirmed resistance (mortality rate <90%) (Fig 2) [16].

In Phase II of the study, the possibly resistant sand fly populations observed during Phase I were reconfirmed for their resistance status. In such repeat bioassays, mortality rate of <98% was interpreted as confirmed resistance in line with WHO test procedures [16]. The populations with confirmed resistance in both Phase I and II were subjected to three additional tests (Fig 2), as follows:

a. The pyrethroid impregnated papers with 5× discriminating concentrations were used to determine resistance intensity.
b. The synergist-insecticide bioassays were performed using PBO 4% impregnated papers in which one set of tests comprised of at least four types of exposures and a batch of 20–25 sand flies for each exposure group; (i) PBO alone – the sand flies were exposed to PBO 4% impregnated paper for 60 minutes, (ii) PBO followed by insecticide – the sand flies were first exposed to PBO 4% impregnated papers for 60 minutes in tubes and then exposed to papers impregnated with either alpha-cypermethrin 0.05%, deltamethrin 0.05%, or lambda-cyhalothrin 0.05% for another 60 minutes, (iii) insecticide alone – the sand flies were exposed to alpha-cypermethrin 0.05%, deltamethrin 0.05%, or lambda-cyhalothrin 0.05% impregnated papers for 60 minutes, and (iv) control – the sand flies were exposed to silicone oil impregnated paper for 60 minutes. A test was considered invalid in case of >10% mortality in the ‘PBO alone’ exposure tube. The interpretation of the synergist-insecticide bioassays was done as follows: (a) the effect of PBO could not be reliably assessed if the mean mortality in tests with ‘insecticide alone’ papers was ≥90%, (b) a monooxygenase-based resistance mechanism was fully accountable for expression of resistance in the test population if the mean mortality in ‘insecticide alone’ tests was <90% and the mean mortality in the ‘PBO followed by insecticide’ tests was ≥98%, and (c) a monooxygenase-based resistance mechanism was partially accountable for the expression of resistance if the mean mortality in the ‘insecticide alone’ tests was <90% and the mean mortality in the ‘PBO followed by insecticide’ tests was more than the mean mortality in the ‘insecticide alone’ tests but <98%.
c. Bioassays with DDT 4% papers along with control were conducted to detect possible cross-resistance to pyrethroids. Risella oil impregnated papers were used in control. The procedures, numbers of replicates and interpretation of mortality percentages of resistance intensity and DDT bioassays were the same as those followed in Phase I.

### Data management and statistical analysis

The number of sand flies found dead in all the test tubes was counted for each replicate and the mortality rate was expressed as a percentage of the dead among the total number of the exposed sand flies. A similar calculation was done for the control tubes. In case the mortality rate in the control tubes was between 5% to <20%, the mortality rate in the test replicates was corrected using Abbott’s formula [20]. Corrected mortality rates were interpreted to determine susceptibility status as described in the WHO guidelines [16] (Fig 2). The tests were discarded if the mortality rate in the control tube was ≥20%.

R software version 4.1.0 [21] was used to conduct all statistical analyses and plotting bar graphs with their mean and standard errors representing variations between the test replicates. For the main statistical analysis of the sand fly mortality, a Poisson regression model was chosen with the number of dead sand flies as the outcome and the total number of the sand flies included in the tests as the exposure. To account for clustering of observations that were done in the same district, or the same village, or that were part of the same susceptibility test, we opted for a Generalized Linear Mixed Model (GLMM) containing random effects for these three levels. In practice, this yielded boundary estimates for the variances of the random effects, which casted doubt on the stability of the model. To better account for the uncertainty of these estimates, we used a Bayesian approach for this analysis using the R package “rstanarm” [22, 23] with default, weakly informative priors. Central credible intervals at 95% were reported for each coefficient and these were interpreted based on the weight of the evidence rather than whether they exceeded 1. Additionally, a probit analysis [24–26] was performed on the R package “ecotox” [27] to estimate the time and fiducial confidence limits at which 50% of the sand fly population were expected to be knocked down (KDT_50_) after exposure to the insecticide impregnated papers.

### Ethical considerations

Ethical clearance for the study was obtained from the Institutional Ethical Review Board of BP Koirala Institute of Health Sciences, Dharan, Nepal. Informed consent was obtained from the household and cattle shed owners where sand flies were collected.

## Results

Of 10,564 female *P. argentipes* collected from the study villages to perform the bioassays during both phases, 8,009 were used in the test replicates and the remaining in the control. Most of the sand flies were blood-fed (>60%), followed by gravid and unfed physiological stages (S1 Fig).

### Phase 1 bioassays

In the IRS villages, the sand fly populations showed resistance to alpha-cypermethrin 0.05% in four villages and possible resistance in one village, with mortality rates between 69.28% ± 3.82% and 92.77% ± 3.01%. The sand flies from four villages showed possible resistance to deltamethrin 0.05% with mortality rates between 95.44% ± 0.12% to 97.95% ± 1.26% and lambda-cyhalothrin 0.05% with mortality rates between 90.12% ± 4.73% to 95.70% ± 2.16%, while full susceptibility was registered in the other remaining villages. The sand fly populations were fully susceptible to bendiocarb 0.1% and malathion 5% (mortality rate >99%) in all the five IRS villages (Fig 3).

**Fig 3.**
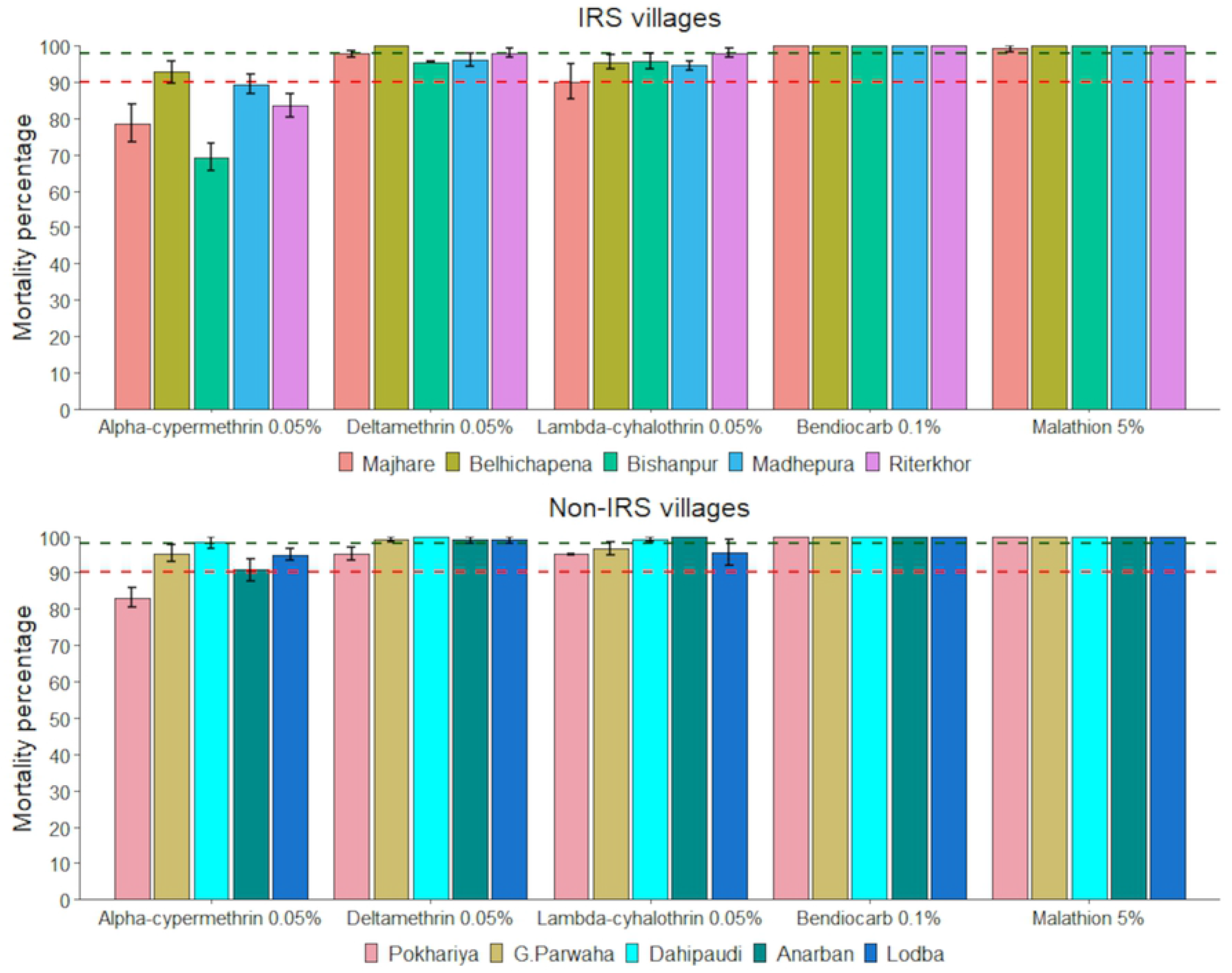
Mean mortality percentage after 24h of holding in the IRS (top) and the non-IRS villages (bottom). Error bars represent the standard error of the mean. The green-dashed intercept represents the level of 98% mortality; the red dashed intercept represents the level of 90% mortality.

In the non-IRS villages, the sand fly populations were resistant to alpha-cypermethrin 0.05% with mortality rate 83.08% ± 2.66% in one village, possibly resistant in three villages with mortality rates between 90.77% ± 3.07% to 95.36% ± 2.24%, and susceptible in one village. All villages except one had vector populations susceptible to deltamethrin 0.05%. In three villages, the sand fly populations showed possible resistance to lambda-cyhalothrin 0.05% with mortality rates 95.04% ± 0.09% to 96.69% ± 1.72%, whereas in two others they were susceptible. Like the IRS villages, the sand flies in all non-IRS villages were susceptible to bendiocarb 0.1% and malathion 5%, with a 100% mortality rate.

Mortality rates were further assessed in the regression model. Overall, mortality in the sand flies was lower in the IRS villages than in the non-IRS villages (IRR = 0.9). While taking account of physiological stages, blood fed and gravid vector sand flies had slightly higher mortalities than unfed ones (IRR >1). Mortality rates were higher (IRR >1) for other insecticides viz. deltamethrin, lambda-cyhalothrin, bendiocarb and malathion than for alpha-cypermethrin. The difference between alpha-cypermethrin and other insecticides in terms of mortality was larger in the sprayed villages than in the unsprayed ones (interaction effect IRR >1). There was little variation between the districts and between the villages, which was not already explained by variables in the model (standard deviations of random effects near 0) (Table 1).

**Table 1.** Bayesian Poisson regression of the sand fly mortality rate in previously sprayed villages, sand fly subtype, and insecticide used in the susceptibility test, with interaction effects between village spraying and insecticide used, and 95% central credible intervals.

|  |  | Incidence rate ratio |
| --- | --- | --- |
| Intercept (base incidence rate) |  | 0.9 (0.8 – 1.02) |
| Village type (ref: Unsprayed) | Village previously sprayed with pyrethroids | 0.9 (0.8 – 1.00) |
| Sand fly physiological status (ref: unfed) | Blood-fed | 1.03 (0.93 – 1.14) |
|  | Gravid | 1.01 (0.91 – 1.13) |
| Insecticide used in test (ref: Alpha-cypermethrin 0.05%) | Deltamethrin 0.05% | 1.07 (0.96 – 1.18) |
|  | Lambda-cyhalothrin 0.05% | 1.05 (0.95 – 1.16) |
|  | Bendiocarb 0.1% | 1.08 (0.98 – 1.20) |
|  | Malathion 5% | 1.08 (0.98 – 1.20) |
| Interaction effect: Village type x Insecticide used in test (ref: Unsprayed x Alpha-cypermethrin 0.05%) | Sprayed x Deltamethrin 0.05% | 1.1 (0.96 – 1.27) |
|  | Sprayed x Lambda-cyhalothrin 0.05% | 1.09 (0.94 – 1.25) |
|  | Sprayed x Bendiocarb 0.1% | 1.12 (0.97 – 1.29) |
|  | Sprayed x Malathion 5% | 1.11 (0.97 – 1.29) |
| Standard deviations of random effects | District | 0.0 (0.0 – 0.0) |
|  | Village | 0.0 (0.0 – 0.0) |
|  | Test | 0.0 (0.0 – 0.0) |

### Proportion of not knocked down and KDT_50_

There was no significant difference in the proportion of sand flies that were not knocked down after 1h exposure with insecticides in the IRS and the non-IRS villages (Fig 4). The estimated time for knockdown of 50% of sand flies (KDT_50_) upon exposure to pyrethroids at the discriminating concentrations (1×) varied between 31.27 to 38.64 min for the IRS villages and 29.73 to 34.86 min for the non-IRS villages. The KDT_50_ was four times shorter (8.70–11.27 min) at 5× concentration of pyrethroids. Higher KDT_50_ (45.27 min) was observed in tests with DDT 4% (Table 2).

**Fig 4.**
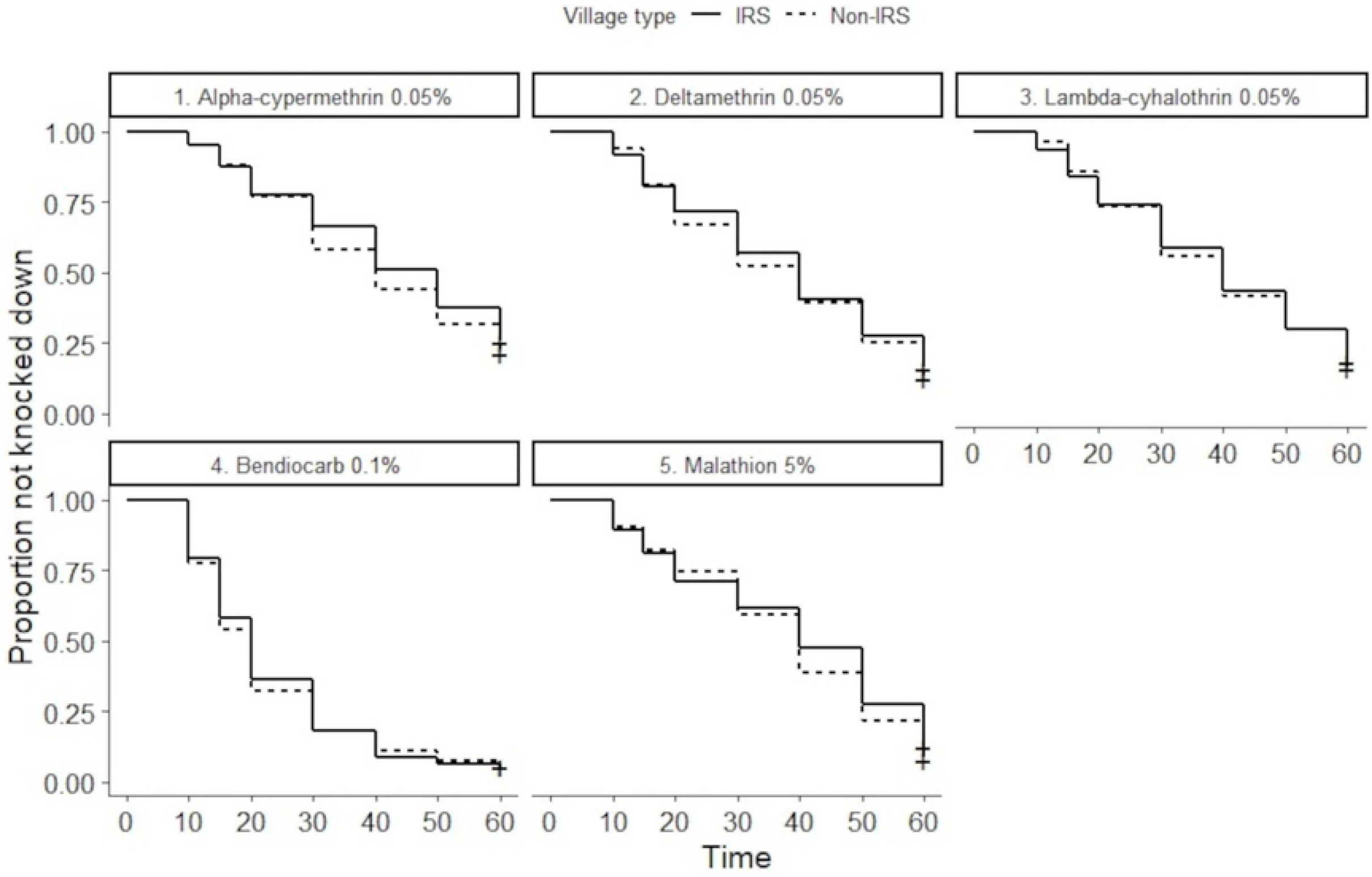
Time until knockdown of *P. argentipes* with different insecticides tested in Phase I in the IRS and the non-IRS villages.

**Table 2.**
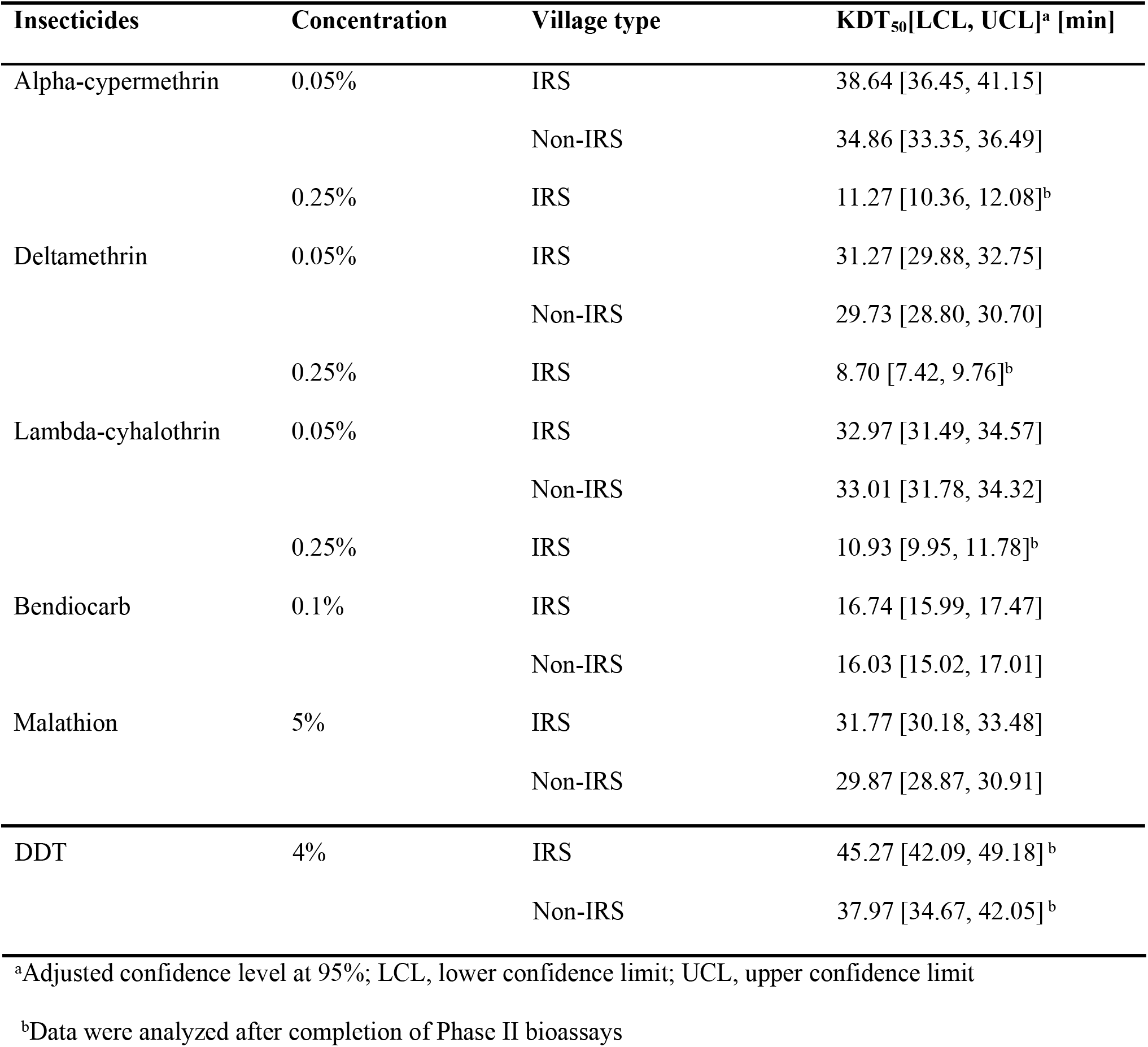
Estimated 50% knockdown times of wild populations of *P. argentipes* sand flies exposed to various insecticide papers impregnated at the 1× and the 5× discriminating concentrations in the IRS and the non-IRS villages.

| Insecticides | Concentration | Village type | KDT<sub>50</sub>[LCL, UCL]<sup>a</sup> [min] |
| --- | --- | --- | --- |
| Alpha-cypermethrin | 0.05% | IRS | 38.64 [36.45, 41.15] |
|  |  | Non-IRS | 34.86 [33.35, 36.49] |
|  | 0.25% | IRS | 11.27 [10.36, 12.08]<sup>b</sup> |
| Deltamethrin | 0.05% | IRS | 31.27 [29.88, 32.75] |
|  |  | Non-IRS | 29.73 [28.80, 30.70] |
|  | 0.25% | IRS | 8.70 [7.42, 9.76]<sup>b</sup> |
| Lambda-cyhalothrin | 0.05% | IRS | 32.97 [31.49, 34.57] |
|  |  | Non-IRS | 33.01 [31.78, 34.32] |
|  | 0.25% | IRS | 10.93 [9.95, 11.78]<sup>b</sup> |
| Bendiocarb | 0.1% | IRS | 16.74 [15.99, 17.47] |
|  |  | Non-IRS | 16.03 [15.02, 17.01] |
| Malathion | 5% | IRS | 31.77 [30.18, 33.48] |
|  |  | Non-IRS | 29.87 [28.87, 30.91] |
| DDT | 4% | IRS | 45.27 [42.09, 49.18]<sup>b</sup> |
|  |  | Non-IRS | 37.97 [34.67, 42.05]<sup>b</sup> |

<sup>a</sup>Adjusted confidence level at 95%; LCL, lower confidence limit; UCL, upper confidence limit
<sup>b</sup>Data were analyzed after completion of Phase II bioassays

### Phase II bioassays

The bioassays were repeated in the second phase in the villages where sand flies were found to be possibly resistant to pyrethroids during Phase I. In all these villages, the presence of resistant phenotypes was confirmed, with mortality rates in all repeated bioassays below 98% (Table 3).

**Table 3.** Assessment of mean mortality percentage of *P. argentipes* to discriminating concentrations of pyrethroids in bioassays in selected villages with possible resistant vector populations detected in Phase I.

| District | Village | Type of village | Insecticide | Mean mortality % (± SE) in Phase I bioassays | Mean mortality % (± SE) in Phase II bioassays |
| --- | --- | --- | --- | --- | --- |
| Saptari | Belhichapena | IRS | Alpha-cypermethrin 0.05% | 92.77 (± 3.09) | 94.98 (± 2.06) |
|  |  |  | Lambda-cyhalothrin 0.05% | 95.44 (± 1.99) | 92.49 (± 1.09) |
| Siraha | Bishanpur | IRS | Deltamethrin 0.05% | 95.44 (± 0.12) | 90.19 (± 2.68) |
|  |  |  | Lambda-cyhalothrin 0.05% | 95.70 (± 2.16) | 94.12 (± 0.98) |
| Sarlahi | Riterkhor | IRS | Deltamethrin 0.05% | 97.95 (± 1.26) | 95.23 (± 2.09) |
|  |  |  | Lambda-cyhalothrin 0.05% | 98.00 (± 1.22) | 96.36 (± 0.91) |
| Sarlahi | Lodba | Non-IRS | Alpha-cypermethrin 0.05% | 94.94 (± 1.58) | 96.13 (± 2.11) |
|  |  |  | Lambda-cyhalothrin 0.05% | 95.45 (± 3.52) | 94.49 (± 1.70) |

SE, standard error of the mean

### Resistance intensity bioassays

The vector population from Bishanpur with confirmed resistance at 1× concentration of all three pyrethroids showed 100% mortality at 5× of the discriminating concentration (S1 Table). As mortality rate at 5× concentration was 100%, bioassays at 10× concentration were not required.

### Synergist bioassays

Mortality rate of the sand flies that were pre-exposed to PBO 4% was higher than of those exposed to a given insecticide only. However, the synergistic effect of PBO to alpha-cypermethrin 0.05% could not be interpreted for the sand fly population in Belhichapena and to lambda-cyhalothrin in Bishanpur as the mean mortality rate in the tests with ‘insecticide alone’ was ≥90%. The susceptibility level was partially restored in Bishanpur and Riterkhor populations for alpha-cypermethrin 0.05%. Interestingly, a complete restoration of susceptibility was seen in Bishanpur population for deltamethrin 0.05% (Fig 5).

**Fig 5.**
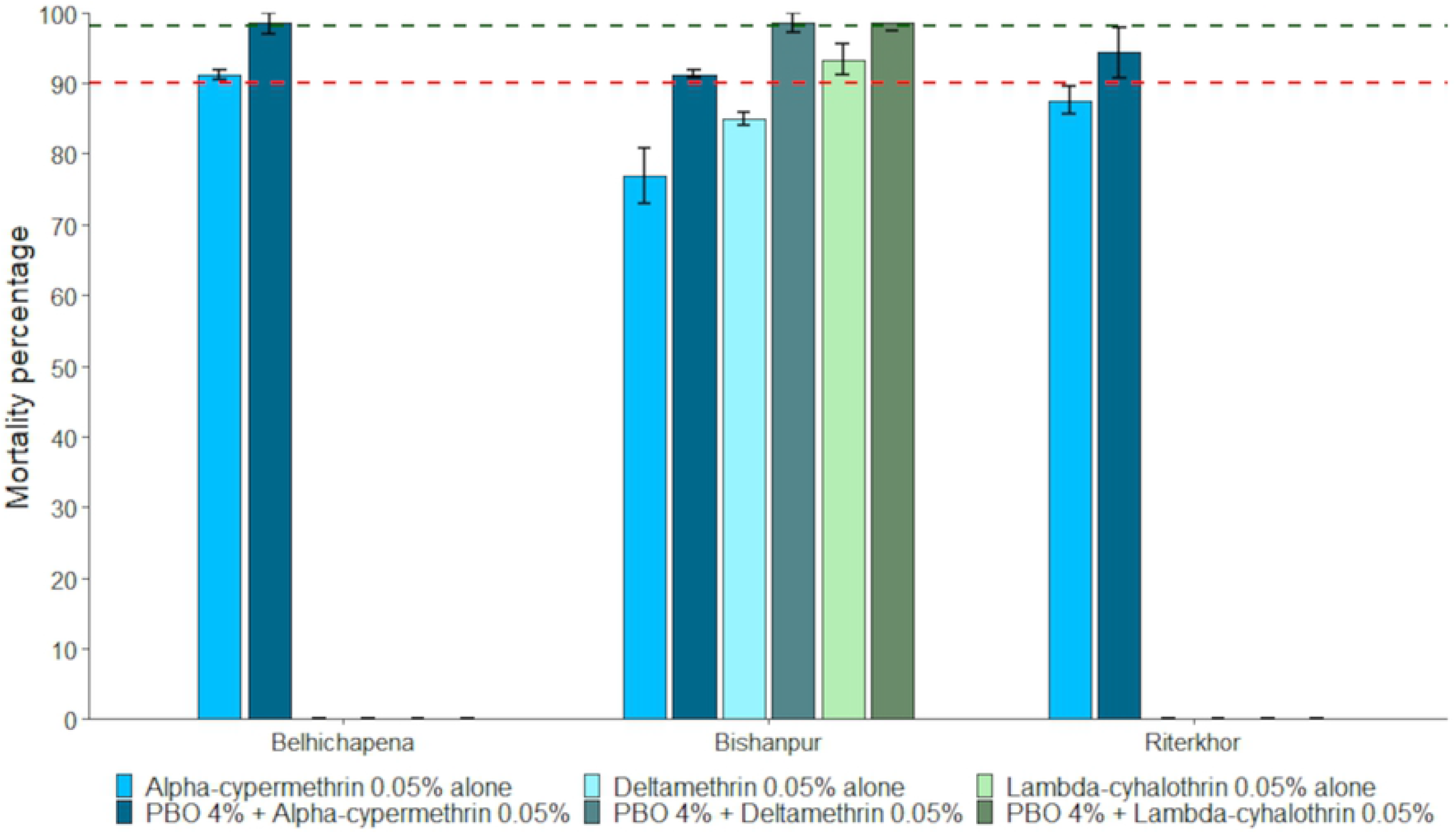
Comparative mortality percentage of *P. argentipes* to pyrethroid insecticides alone or pre-exposed to PBO 4% papers followed by pyrethroid insecticides. The error bars represent the standard error of the mean. The green-dashed intercept represents the level of 98% mortality; the red dashed intercept represents the level of 90% mortality.

### Bioassays with DDT 4%

Mortality rate in the pyrethroid resistant *P. argentipes* populations exposed to DDT 4% papers ranged between 55.14% ± 1.49% and 76.15% ± 3.51%. Mortality percentage of pyrethroids in the corresponding villages were included from Phase I for the assessment (Fig 6).

**Fig 6.**
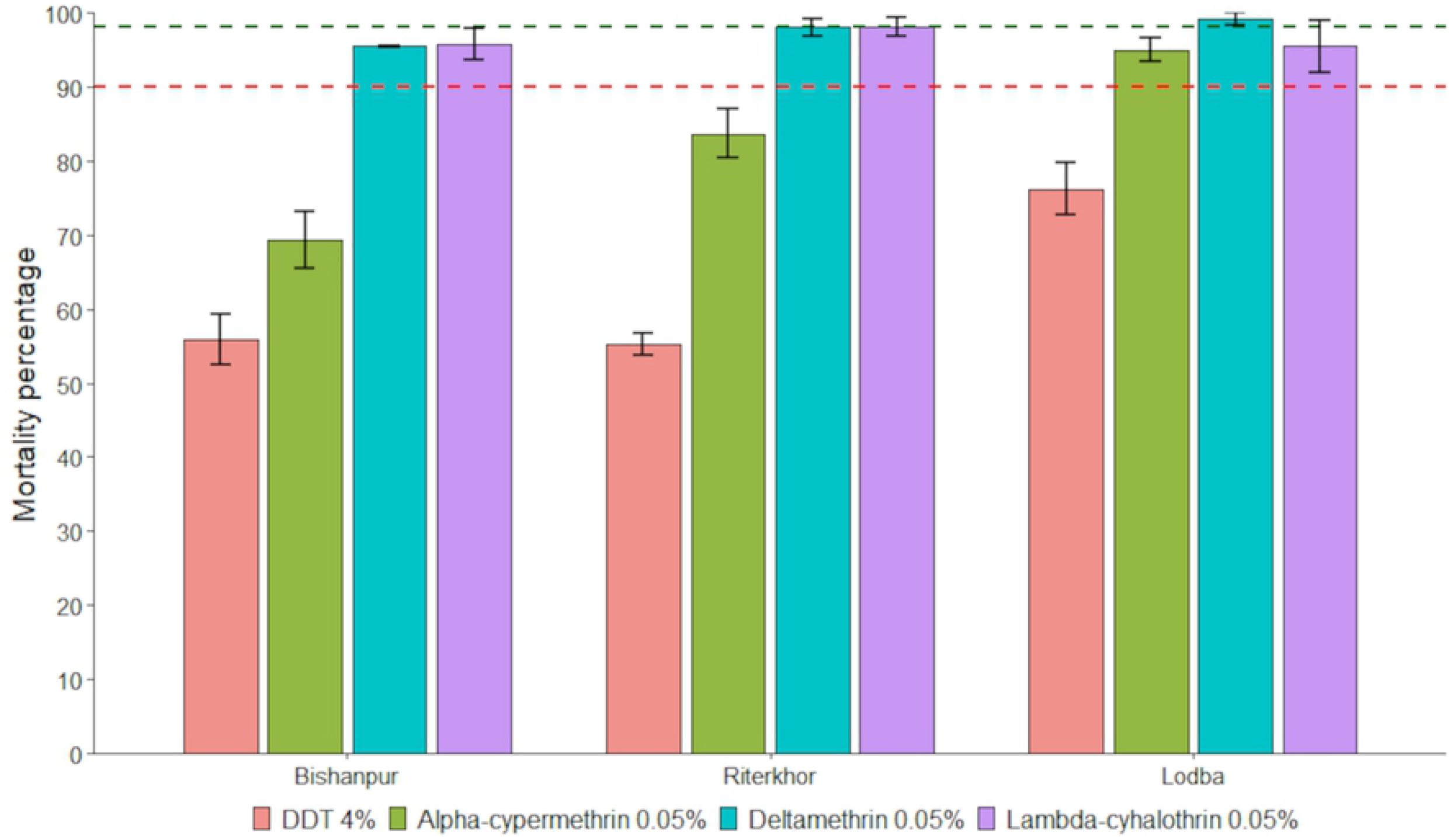
DDT-pyrethroid cross-resistance in *P. argentipes* from the selected IRS and non-IRS villages. The error bars represent the standard error of the mean. The green-dashed intercept represents the level of 98% mortality; the red dashed intercept represents the level of 90% mortality.

## Discussion

In our study, we found that pyrethroid resistance in the *P. argentipes* populations is well established in all the IRS villages, particularly so for alpha-cypermethrin 0.05% which has already been used for more than 15 years now in the IRS campaigns in Nepal. Moreover, pyrethroid resistance was also seen in the non-IRS villages. Development of pyrethroid resistance in the vector populations was also supported by a similar KDT_50_ in both the IRS and the non-IRS villages. Nevertheless, vector populations in all the study villages were fully susceptible to the selected carbamate and organophosphate insecticides. The resistance intensity bioassays confirmed the intensity of pyrethroid resistance is low, suggesting that a possible operational failure of pyrethroid-based vector control is unlikely in the immediate future, although it signals to implement an insecticide resistance management approach. Incomplete restoration of susceptibility in PBO synergist bioassays was indicative of a partial involvement of monooxygenase-based metabolic resistance mechanism. Resistance to DDT in pyrethroid-resistant populations is indicative of presence of *kdr* mutations in the target site gene (*vgsc*) of the vector sand flies. As bioassays were conducted using wild captured sand flies, blood-fed stages outnumbered gravid and unfed in both categories of the villages (IRS and non-IRS), exposures (test and control) and outcomes (dead and alive) with expected mortalities slightly higher in blood fed than the other two physiological stages.

Monitoring of the susceptibility of wild populations of *P. argentipes* is essential to sustain the effectiveness of insecticides, as this can guide insecticide resistance management. While sporadic information on the susceptibility status of *P. argentipes* in Nepal were available over the last 25 years, resistance management has not been pursued with priority within the national VL vector control programme framework. The first known investigation in Nepal, conducted in Thilla village of Dhanusha district, dates back to 1997 and reported that *P. argentipes* populations were susceptible to lambda-cyhalothrin 0.1% and deltamethrin 0.25% [28]. In subsequent publications until 2001, the *P. argentipes* populations were reported to be susceptible to DDT 4%, permethrin 0.25%, deltamethrin 0.25%, lambda-cyhalothrin 0.1%, malathion 5% and bendiocarb 0.1% [28, 29]. In 2009, bioassays conducted in four villages from Eastern Nepal revealed the presence of DDT resistance in one of the four populations of *P. argentipes*, as well as possible resistance to deltamethrin 0.05%, thereby providing an early sign of development of selection pressure against pyrethroids [10]. Bioassays in three districts of Eastern Nepal in 2015 and 2016, on the other hand showed susceptible vector populations at 60 min exposure to alpha-cypermethrin 0.05% and deltamethrin 0.05% [12], which is contrary to the results obtained in our study.

Most of the VL endemic districts in Nepal share close borders with VL endemic districts of Bihar state in India, where DDT based IRS was replaced with a pyrethroid in 2015 due to growing evidence of DDT resistance in vector sand flies [11]. A similar situation was experienced in the bordering part of Nepal in a previous study [10] as well as in our current study. Since then, several reports have been published reporting phenotypic resistance to pyrethroids [30–33]. Recently, the presence of *kdr* mutations in vector populations in some parts of India hinted the emergence of pyrethroid resistance at molecular level [11, 30]. Although *P. argentipes* populations were found susceptible to malathion in several locations within South-East Asia [30, 31, 34, 35], resistant populations have also been detected [35]. In a recent study in three states of India, *P. argentipes* populations showed full susceptibility to all insecticides except for DDT [36]. In Bangladesh, another VL endemic country neighboring to Nepal, the *P. argentipes* populations have shown to be susceptible to insecticides in major classes used in IRS till date [12]. Only a few hundred kilometers apart, information on the susceptibility status of *P. argentipes* in VL endemic areas in India and Bangladesh gives insight into the necessity of such investigations in Nepal.

The delayed knockdown time and reduced mortality detected in our study illustrate the onset of pyrethroid resistance in vector populations. Both the IRS and the non-IRS villages showed similar knockdown times in vector populations, suggesting the spread of pyrethroid resistance in endemic areas irrespective of IRS interventions in recent years. Other causes contributing to the development of resistance in this region might be the widespread use of agricultural pesticides. Unfortunately, no baseline data on KDT for susceptible reference strains of *P. argentipes* are available in South-East Asia to compare our findings with. The lethal time estimated for 50% mortality (LT_50_) of the *P. argentipes* colony population of Rajendra Memorial Research Institute of Medical Sciences, India was <15 min for deltamethrin 0.05% [10], almost half of the time predicted for knockdown in our study. Another study from India [30] reported similar knockdown times as observed in our study (KDT_50_) for wild caught *P. argentipes* against different classes of insecticides. The fact that the highest KDT_50_ was observed for DDT 4% was not surprising, as resistance in the vector population for this insecticide was already well-established and acknowledged.

One of the major strengths of this study is the baseline information generated on the susceptibility status of the alternative classes of insecticides, carbamate and organophosphate, which will be valuable for evidence-based decision making in insecticide resistance management of sand flies. In addition, the observed low intensity of resistance to pyrethroids suggests that these insecticides are not completely obsolete and supports the possibility of reversal to susceptibility in vector sand flies provided that alternative insecticides are used for a few years as resistance mitigation measures. Also, for the first time, we attempted PBO synergist-pyrethroid bioassays to explore possible underlying monooxygenase-based metabolic resistance mechanism. Presence of pyrethroid-DDT cross-resistance can be assessed as a proxy indicator of presence of *kdr* mutations within the *vgsc* gene in nerve cells for the expression of resistance [11, 30]. In the context of confirmed resistance against pyrethroids, if the VL control programme in Nepal plans to switch to alternative classes of insecticides for IRS, organophosphates could be a choice considering the monooxygenase-based metabolic cross-resistance between pyrethroid and carbamate classes of insecticides [37].

The procedures for conventional bioassays are sophisticated in terms of resources and skills, however, these are WHO approved and provide strong evidence on the focal status of phenotypic susceptibility of a particular vector species to a specific insecticide. One of the limitations of this study is that we had to implement the methodology and insecticide discriminating concentrations meant for testing the susceptibility of mosquitoes due to the absence of WHO standardized procedures and specific insecticide discriminating concentrations for testing the susceptibility of sand flies. Nonetheless, several authors have preceded us in the use of this adopted methodology to assess the susceptibility status of sand flies in the Indian subcontinent [10, 12, 30, 36]. A recent study in Sri Lanka has quantified far lower lethal concentrations of insecticides to kill 100% (LC_100_) of *P. argentipes* as compared to discriminating concentrations used to monitor resistant status of mosquitoes [38]. In that case the resistance status observed in our study would likely be an underestimation. To overcome this limitation, WHO is undertaking a global multi-centre study for determining insecticide discriminating concentrations of various insecticides for sand flies and the results are expected in 2022 (RS Yadav, personal communication). Another limitation of the study was that we could not standardize the age and physiological stages as we conducted the bioassays with wild captured adult sand flies. On the other hand, the fact that the age distribution and physiological stages of the sand flies caught were representative of the real-time wild vector population [16], this could be considered as a strength of the study as well.

We cannot assume that the susceptibility status of the vector sand flies observed in the five selected VL endemic districts of Eastern and Central Nepal are representative for other endemic districts from Western part or the hilly districts located in diverse geo-ecological zones and having unlike IRS history. Monitoring of the susceptibility status should therefore be carried out in diverse geo-ecological areas, to allow for area-specific recommendations. Non-endemic districts should not be excluded from this exercise, as sporadic VL outbreaks might warrant effective evidence based vector control interventions as well.

## Conclusion

After three decades of IRS with a single class of insecticides in Nepal, we confirmed the presence of pyrethroid resistance in *P. argentipes* sand flies. It is strongly recommended that the national vector-borne disease control programme takes this evidence into account when deciding on insecticides to be used for future IRS activities, and switches to an alternative class of insecticides with a different mode of action such as organophosphates or novel products prequalified by WHO. Also, this study endorses the need for regular and systematic monitoring of the susceptibility status of the vector of VL to inform evidence-based decision making for the use of insecticides in the fight against VL.

## Acknowledgements

We would like to thank all the female community health volunteers from the respective study villages for guiding us through the village and facilitating communication with the community people. We are also grateful to the vector control officers from the respective district public health offices. We appreciate members of our field team; Manish Karn, Kailash Majhi, Jibachh Sah and Durga Hasda for their sincere effort for collection of adequate sand flies and to assist in conduction of bioassays.

## Author Contribution

**Conceptualization:** LR, SU, KC, WVB

**Data curation:** LR

**Formal analysis:** LR, TS

**Funding acquisition:** LR, SU

**Methodology:** LR, SU, KC, UK, URP, MLD, RSY, WVB

**Project administration:** LR, SU, KC

**Resources:** LR, SU, KC

**Software:** LR, TS

**Supervision:** KC, UK, URP, MLD, WVB

**Validation:** MLD, RSY, WVB

**Visualization:** LR

**Writing — original draft:** LR

**Writing — review & editing:** LR, SU, KC, TS, UK, URP, MLD, RSY, WVB

## Supporting information

**S1 Fig. Percentage composition of physiological stages of female *P. argentipes* used in the test (T) and the control (C) replicates.** Inside the doughnut chart: A, alive and D, dead sand flies after 24h recovery period. UF, unfed; BF, blood-fed and G, gravid *P. argentipes*.

**S1 Table. Assessment of mean mortality percentage of *P. argentipes* to 5× discriminating concentrations of pyrethroids in a selected village with resistant sand fly populations in Phase I bioassays.**

**S1 Data. Dataset of sand flies collected, insecticide tested and results of bioassays in detail.**

